# Directionally Biased Neuronal Responses to Pitch-Axis Vestibular Stimulation in Larval Zebrafish Compared to Roll-Axis Responses

**DOI:** 10.1101/2024.03.22.586054

**Authors:** Geoffrey Migault, Natalia Beiza-Canelo, Sharbatanu Chatterjee, Georges Debrégeas, Volker Bormuth

## Abstract

The vestibular apparatus plays a pivotal role in maintaining postural equilibrium and processing movement signals, underscoring its importance in studying certain neurological disorders. Here, we present a rotating light-sheet microscope, enabling brain-wide functional recordings during dynamic vestibular stimulation along the pitch axes in addition to the roll axis in head-restrained zebrafish larvae. The system incorporates a double galvanometer mirror configuration, is amenable to 3D printing, and enhances scanning efficiency. Employing this apparatus, we have successfully conducted the first comprehensive mapping of zebrafish brain responses to dynamic pitch-tilt vestibular stimulation. Through Fourier and regression analyses, we report an asymmetry in neuronal recruitment during nose-up versus nose-down pitch tilts within critical regions, including the cerebellum, oculomotor nucleus, caudal hindbrain, and vestibular nucleus, highlighting physiological adaptations to downward motion. We identified specific brain regions, notably the cerebellum and medial-rostral rhombencephalon, that respond to roll-but not pitch-tilt vestibular stimuli. Furthermore, we have identified a transgenic line that closely correlates with our functional mappings and demonstrates a significant response to vestibular stimulation. The elucidation of brain-wide neuronal circuits involved in vestibular processing establishes a foundational framework for subsequent detailed investigations into the molecular and genetic mechanisms underlying postural control and motion perception.

## Introduction

The vestibular apparatus is a critical sensory system in animals, essential for maintaining postural control, perceiving spatial orientation, and responding to movement cues. It plays a fundamental role in coordinating motor behaviors, crucial for balance and spatial awareness. Dysfunction in this system is associated with various neurological disorders such as Alzheimer’s disease, Parkinson’s disease, progressive supranuclear palsy, and autism spectrum disorder [1, 2, 3, 4], underscoring the importance of understanding its mechanisms at the brain-wide circuit level. Recent advances in imaging techniques have provided opportunities to explore brain-wide neural activity related to vestibular processing. For instance, a miniaturized rotating light-sheet microscopy approach [5] allows comprehensive brain-wide neuronal recordings during controlled vestibular stimulation in larval zebrafish. Other methods include tilting fish under a static microscope [6], rotating them with the microscope objective [7, 8], and creating fictive vestibular stimuli by exerting forces on the otoliths with optical tweezers [9] or utilizing magnets following ferrofluid injection into the otic vesicle [10]. Comprehensive brain wide neural responses have only been characterized for the roll-tilt axis. In contrast, investigations of pitch-tilt responses have been confined to distinct sparse neural subpopulations [11, 6], leaving the broader, pan-neuronal correlates associated with pitch-tilt stimulation unexplored.

This study introduces significant enhancements to the miniaturized rotating light-sheet microscope approach, enabling precise rotations along axes between roll and pitch tilt directions with simultaneous whole-brain imaging. Our novel design, which is 3D printable and easy to assemble and reproduce, features a double galvanometer mirror configuration, enhancing scanning capabilities. This advancement allows us to examine larval zebrafish responses to vestibular stimulation on the pitch-tilt axis and compare them to the roll-tilt response, an aspect of vestibular processing previously unexplored with whole-brain imaging.

Maintaining postural control along the pitch-tilt axis is critical for survival, often closely tied to the initiation of movement. For instance, organisms need to lean forward to start moving, causing disbalance, which needs to be compensated. This balancing act, especially against acceleration and deceleration forces, is crucial. Larval zebrafish present a unique case in this context. As demonstrated by Ehrlich et al. [12] they experience inherent postural instability along the pitch axis, attributed to the misalignment between their center of buoyancy and center of gravity. This misalignment leads to a consistent nose-down postural destabilization. To counteract this, larval zebrafish perform active righting movements, accompanied by changes to pitch angle to maintain their posture. Notably, the initiation of their characteristic swim behavior patterns is directly influenced by their vestibular system’s detection of this destabilization. This highlights the evolutionary necessity for developing efficient neuronal pathways that compensate for pitch-axis postural control and its related movement complexities.

Our primary objective was to determine if distinct neuronal responses and potential asymmetries exist in processing nose-up and nose-down pitch-tilt directions, given the behavioural ubiquity of the nose-up pitch-tilt direction movements used in righting movements. We also recorded responses to roll-tilt stimuli in the same animals to compare the spatial organization of neuronal populations between pitch-tilt and roll-tilt axes. This offers insights into how different vestibular stimulations differentially activate brain regions.

## Results

### Enhanced Light-Sheet Microscopy for Diverse Vestibular Stimulation

The light-sheet unit is an advanced version of the setup detailed by Migault et al. [5]. Modifications include a 3D printable bracket and a dual galvanometer mirror setup. The system utilizes a 488 nm laser beam delivered via an optical fiber. This beam, first collimated by a 5x objective (numerical aperture 0.16), is directed to the first galvanometer mirror, enabling z-directional light sheet translation. The beam then reaches the second galvanometer mirror, oscillating at 400 Hz, creating a light sheet parallel to the imaging plane. The core of the unit is a custom-designed bracket that secures the objectives and mirrors. This arrangement is shown in Figure 1. The 3D design file is open source and can be interactively inspected at https://a360.co/2U1KBoH, or accessed via its persistent DOI: 10.5281/zenodo.15498761 [13]. For ease of adjustment, the objectives are clamped rather than screwed, which ensures precise positioning of the optical fiber output and the laser beam for optimal collimation. Additionally, the illumination objective’s translation feature simplifies the alignment of the laser beam’s waist with the observation objective’s field of view center. We reinforced the 3D-printed unit with a thin aluminum plate on one side to achieve mechanical stability. This modification ensures an ultra-stable setup, limiting movement in the x and y directions to less than 2 *µ*m and in the z-direction to less than 500 nm, as depicted in Figure 1e.

**Figure 1:**
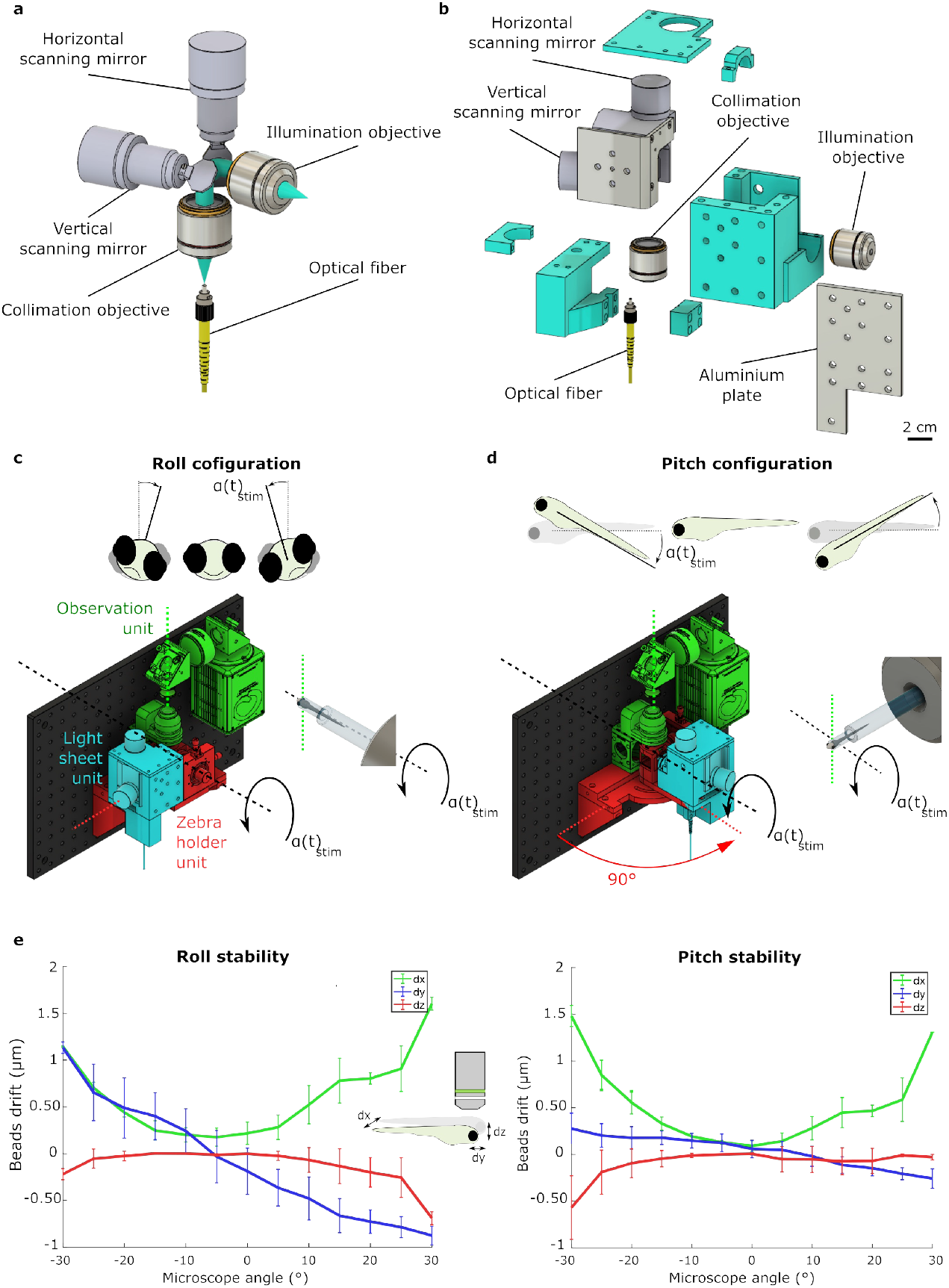
The Experimental Setup. a) Schematic representation of the optical elements that define the laser path in the light-sheet unit, with the laser path highlighted in blue. b) Exploded view of the light-sheet unit, displaying the 3D printed components in cyan. The metal plate (gray) is securely attached to the 3D-printed bracket to enhance the mechanical stability of the unit during microscope rotations. c) Full view of the light-sheet microscope setup: The light-sheet unit is depicted in blue, the specimen holder in red, and the detection unit in green. These components are mounted on a breadboard fixed to a high-speed precision rotation stage. The light-sheet unit and the sample holder were placed on a small rotatable platform to allow arbitrary changes between the roll-tilt and pitch-tilt configurations for vestibular stimulation. The position of this element was adjusted to define the axis around which the fish was tilted when the entire microscope was rotated. The configuration displayed here demonstrates a vestibular roll-tilt stimulus. d) The configuration displayed here demonstrates a vestibular pitch-tilt stimulus. e) Mechanical stability assessment along the three directions (dx, dy, and dz) as a function of the microscope rotation angle. Error bars represent the measurements’ standard deviation (SD), with N = 5. The microscope rotation spans from −30° to +30° for the characterization along the roll- and pitch-tilt axes.

To broaden the scope of vestibular stimulation directions, we mounted both the light-sheet unit and the sample holder on a compact, rotatable platform. This platform’s rotation axis, depicted as a green dotted line in Figure 1c), is aligned with the detection objective’s optical axis. By rotating and then securing this element with a fixation screw, we defined the axis around which the fish tilts when the entire microscope rotates. Consequently, we could switch the vestibular stimulation direction from a roll-tilt (Figure 1c) to a pitch-tilt (Figure 1d) configuration. Crucially, the fish consistently stays within the microscope’s field of view during these manipulations. Also, as the light-sheet unit shares the rotating platform with the sample chamber, it always maintains a lateral orientation relative to the fish. This setup enables us to consistently obtain sharply focused images of the same neurons by merely adjusting the vestibular stimulation axis while keeping the illumination uniform. Such a configuration is instrumental in directly comparing the neuronal responses to roll and pitch-tilt stimulations. The microscope roll and pitch axes are, by design, orthogonal to each other and to the gravity field. In practice, however, the fish cannot always be mounted with perfect alignment relative to the rotation axis. Such misalignment can introduce crosstalk between the roll- and pitch-evoked responses. We quantified the misalignment angles for each experiment and analytically estimated the resulting crosstalk. Across all datasets, the predicted crosstalk remained below 5% on average, making it negligible for the conclusions drawn in this study (see Methods).

### Comparing brain-wide neuronal correlates of pitch- and roll-tilt stimuli

#### Neural Responses Induced by Sinusoidal Pitch-Tilt Stimuli

Utilizing our improved system, we investigated the brain’s neuronal response to pitch-tilt vestibular stimulation. We used the Tg(elavl3:H2b-GCaMP6f) transgenic zebrafish line, which exhibits panneural expression of the GCaMP6f calcium sensor. Animals were subjected to a sinusoidal rotation around their horizontal dorsal up orientation, at a frequency of 0.2 Hz and an amplitude of ±20°, for 10 minutes (120 periods), which corresponds to a peak velocity of 12.5°/s A substantial number of neurons responded to this stimulation, as showcased in Video 1.

We successfully mapped brain-wide neuronal responses using Fourier analysis to pinpoint neurons showing significant activity at the stimulus frequency. The phase of these responses was analyzed relative to the stimulus waveform’s phase, revealing the intricate dynamics of neuronal activity (for details, see Methods: Phase Maps). We corrected for delays caused by the calcium sensor to determine the actual neural response phase (see Methods: Decay Time Constant of Calcium Sensor). Our analysis across the brain revealed a spectrum of phase shifts, peaking around *π*/3 and 4*π*/3, as shown in Figure 2a. To visualize the spatial distribution of neurons based on their response phases, we color-coded them to create phase maps depicted in Figures 2c-e and 3D animated in Video 1. Here, color coding represents phase shift values, and pixel intensity indicates response amplitude.

**Figure 2:**
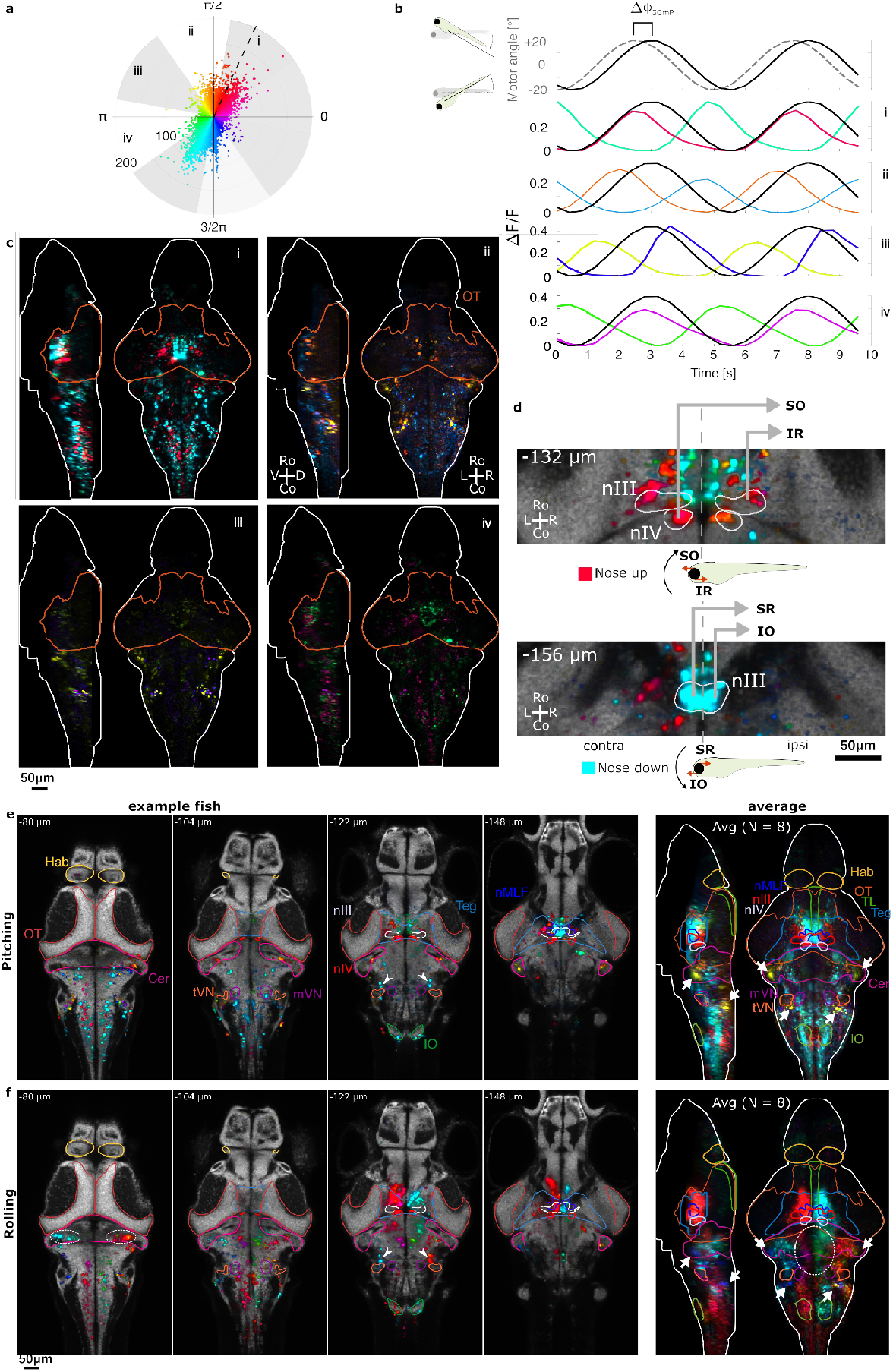
Neuronal Response Maps to Sinusoidal Pitch- and Roll-Tilt Stimulation. a) Polar plot displaying neuron fluorescence response amplitude, measured in signal-to-noise ratio units at the stimulation frequency, plotted against the response phase delay relative to the stimulus. Phase delays were adjusted for the 1.15 radians phase delay introduced by the GCaMP6f sensor (see Methods Decay Time Constant of Calcium Sensor). The dotted line marks the zero phase shift position before correction. Phase delays are color-coded, and this color code is consistent across other panels. b) Top: Microscope rotation angle over time corresponding to the vestibular sine stimulus (dashed line) and phase-shifted to compensate for the GCaMP6f sensor delay (black solid line). Bottom: Neuron-averaged traces for the four phase windows, color-coded according to their respective average phase shifts. c) Maximum projection views of a representative example fish’s phase map, registered to the Z-Brain reference brain (as depicted in (e), divided into four phase intervals (i, ii, iii, and iv) as shown in (a). Each phase interval is displayed as maximum projection from left to midline and from dorsal to ventral. d) Close-up view of the oculomotor (nIII) and trochlear (nIV) nuclei, revealing the spatial organization of their functional responses consistent with the anatomical projection patterns of these motoneurons to extraocular muscles (superior oblique (SO), inferior rectus (IR), superior rectus (SR), inferior oblique (IO)). The top and bottom panels correspond to the nose-up and nose-down positions, respectively, with schematics indicating the direction of stimulation (black arrow) and the direction of force exerted by the extraocular muscles (orange arrows) associated with pitch stimulation. e) Pitch tilt response. Left: Four selected layers of the example fish’s phase map are shown. White arrowheads indicate vestibulospinal neurons. Right: Average phase maps of the pitch-tilt response (N = 8 fish). Arrows highlight neural populations with phasic responses in the lateral cerebellum and dorsal hindbrain. f) Roll tilt response. Left: Four selected layers of the example fish’s phase map are shown, displaying the same neurons as in (e). White arrowheads indicate vestibulospinal neurons. The dashed oval highlights roll-tilt responses in the cerebellum, which are absent in the pitch-tilt maps. Right: Average phase maps of the roll-tilt response (N = 8 fish, same individuals as in (e), right). Arrows highlight the same neural populations with phasic responses in the lateral cerebellum and dorsal hindbrain, as in (e, right). The dashed oval highlights roll-tilt responses in rhombomeres 1–3, which are absent in the pitch-tilt maps. All recordings shown in this figure were obtained using the Tg(elavl3:H2b-GCaMP6f) transgenic zebrafish line. The response maps were registered to the ZBrain atlas [14] and are displayed in this space, overlaid with selected brain region contours extracted from the ZBrain atlas. The highlighted regions include the optic tec tum (OT), torus longitudinalis (TL), habenula (Hab), cerebellum (Cer), tegmentum (Teg), oculomotor nucleus (nIII), trochlear nucleus (nIV), nucleus of the medial longitudinal fasciculus (nMLF), tangential nucleus (tVN), medial nucleus (mVN), and inferior olive (IO). Abbreviations: C: caudal; D: dorsal; L: left; R: right; Ro: rostral. Refer also Video S1, S2 and S3 and Figure S2 and S1.

The response maps exhibited a mirror-symmetrical layout around the sagittal midplane. For enhanced clarity, we displayed the phase map of a typical fish in four separate phase windows using z-projections (Figure 2b), with Figure 2a illustrating how phases were allocated for each window spanning a phase and its direct 180° opposite. Remarkably, within each window, the neuronal populations displaying responses 180° out of phase with each other show distinct and separated spatially organized patterns. We centered the first phase window around *π*/3 and 4*π*/3, where we observed the largest response amplitudes. Figure 2c shows the mean neural activities of these so-defined neuronal clusters, illustrating how they are tuned to different parts of the stimulus cycle. Neurons in the phase window (i), which encompassed most of the neurons, exhibited a phase-shifted response relative to the stimulus (see Figure 2b), indicating sensitivity to both the pitch angle and the angular velocity of the stimulus. In a sinusoidal stimulation paradigm, such a phase shift arises when the neural response depends not only on the stimulus itself but also on its rate of change. A response that combines sensitivity to position and velocity leads to an intermediate phase shift between zero and ninety degrees, as the derivative of a sine function is a cosine, which is phase-shifted by 90 degrees. By contrast, sensitivity to position alone—or to position combined with acceleration—does not result in a net phase shift. The observed shift therefore reflects a mixed sensitivity to both the angular position and velocity of the pitch stimulus. These average responses were tuned to nose-up pitch-tilt (red) and nose-down pitch-tilt (cyan). To further comprehend the three-dimensional organization, we show single-layer views of the phase maps for four selected layers in Figure 2c. Strikingly, the response patterns we found are stereotypical among different animals, as shown by the computed averaged phase maps from recordings in 8 fish (Figure 2e, right).

Localized neuronal activity was observed in the tegmentum, encompassing the nucleus of the medial longitudinal fasciculus (nMLF), oculomotor and trochlear nuclei, as well as in the cerebellum and the rhombencephalon, particularly in the octavo-lateral nucleus regions adjacent to the otic vesicles. This region includes the tangential and medial vestibular nuclei and vestibulo-spinal neurons. Furthermore, the inferior olive and parts of rhombomere 7 were responsive. More in details, in the tegmentum, we found that the trochlear nucleus (nIV) notably responded solely to nose-upward movements. The oculomotor nucleus (nIII) displayed a differentiated response pattern: its dorsal part was activated by nose-up movements, while the ventral part reacted to nose-down movements. This asymmetry in oculomotor neuron activation in nIII and nIV aligns with findings from recent backtracing studies mapping these neurons to their specific oculomotor muscle targets [15]. The functional responses’ spatial organization reflects the projection patterns of these ocular motor neurons, which are pivotal for controlling eye rotations during pitch-tilt movements of the body, either nose-up or nose-down (see Figure 2d and Figure S2). During a nose-down body pitch rotation, the rostral part of both eyes rotates upward while the caudal part rotates downward, ensuring a stable gaze for both eyes. Conversely, during a nose-up body pitch rotation, the rostral part of the eyes rotates downward, stabilizing the visual field. Specifically, during nose-up rotations, the nIV nucleus, targeting exclusively the superior oblique eye muscles, in conjunction with the dorsal part of nIII, which innervates the inferior rectus eye muscles, coordinates downward eye rotations. In contrast, nose-down pitch-tilt movements activate nIII’s ventral part, associated with the superior rectus and inferior oblique eye muscles, facilitating upward eye movements. In addition to these oculomotor regions, we observed a medial-dorsal cluster of neurons in the caudal hindbrain that responded selectively to nose-up movements, flanked by two laterally located strips of neurons tuned to nose-down movements (Figure 2e).

### Neural Responses to Sinusoidal Roll-Tilt Stimuli in the Same Fish

Beyond documenting the pitch-tilt response, we also captured the roll response in all fish in response to a sinusoidal rotation along the roll-tilt axis with an amplitude of ±20° around their preferred dorsal-up posture. Figure 2f and video 3 depict phase maps corresponding to these roll responses. The pitch-tilt and the roll-tilt stimulus recruited globally the same brain regions, including the optic tectum, habenula, cerebellum, tegmentum, oculomotor nucleus nIII, trochlear nucleus nIV, medial longitudinal fasciculus nucleus, tangential nucleus, inferior olive, and rhombomeres Rh1 to Rh7. We observed significant differences between the roll- and pitch-tilt responses (Fig. 2e and Fig. 2f). While both responses are mirror-symmetric to the sagittal midplane, the roll-tilt response shows a distinctive 180-degree phase shift in neuronal responses between the two brain hemispheres. We observed populations of neurons in the cerebellum that responded only to the roll-tilt stimulus (Figure 2f dashed oval compared to e), and in general, fewer cerebellar neurons were recruited during the pitch-tilt stimulus. Additionally, the medial-rostral part of the rhombencephalon exhibited exclusive responsiveness to the roll-tilt stimulus. Vestibulospinal neurons were strongly activated on each brain side during roll- and pitch-tilt stimulation (see arrowheads in Fig. 2e-f). However, during pitch-tilt stimulation, these neurons demonstrated tuning to nose-down rotations without responding to nose-up movement. At the same time, during roll-tilt, they displayed tuning to ipsilateral ear-down rolls, respectively. Furthermore, we observed, during both stimulation directions, the recruitment of a population of neurons in the tegmentum, rostral to the region labeled in the zBrain atlas as nMLF, which could correspond to neurons of the interstitial nucleus of Cajal.

### Neural Responses to Stepwise Pitch-Tilt Stimuli

Alongside sinusoidal stimulation, we analyzed the brain-wide response to step stimuli along the pitch-tilt axis in the same fish. The fish were pitch-tilted in alternating nose-up and nose-down directions from their horizontal position, cycling through three amplitudes (10°, 15°, 20°) as illustrated in Figure 3. Each step transition occurred at a peak angular velocity of 30°/s, followed by a 20-second dwell time at the new position. Between each step, the fish were returned to the horizontal (0°) position for an additional 20-second dwell period. This sequence of six steps (three amplitudes in both directions, each separated by a return to 0°) was repeated four times in total.

**Figure 3:**
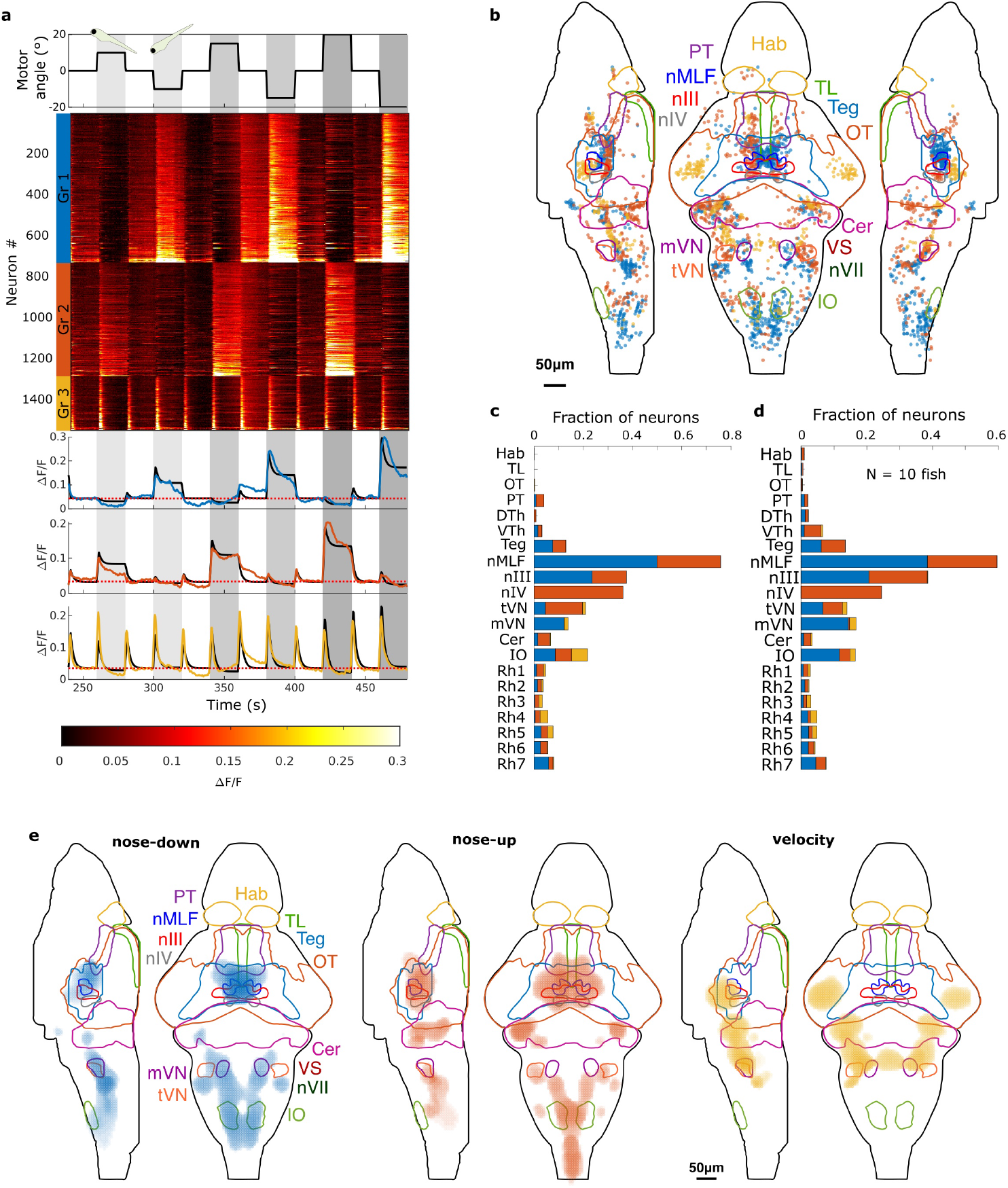
Brain-wide Response to Step-wise Vestibular Stimuli along the Pitch-Tilt Axis. a) Trial-averaged neuronal responses, of an example fish, ordered by group affiliation determined through multilinear regression analysis against tonic and phasic components of the step stimulus. Orange indicates neurons with strong tonic and/or phasic responses to nose-up pitch-tilt; blue indicates neurons with strong tonic and/or phasic responses to nose-down pitch-tilt; yellow indicates neurons with phasic, adirectional responses. Top: Trace of the rotation angle of the stimulus (in black). Positive angles correspond to nose-up, and negative angles correspond to nose-down. Middle: Rasterplot of trial-averaged responses of all identified neurons. Neuron indices are shown on the left, and the color bar indicates group membership. Within each group, neurons are ranked based on the value of the group coefficient. Bottom: Mean trial-averaged response across all neurons of their respective groups, shown in their respective colors. The black line represents the average regression fit. b) Spatial organization in the brain of neurons of the three response groups. c) Histogram showing the percentage of neurons in selected brain regions that belong to the different response groups. d) Similar analysis to (c), but displaying cross-animal results (N = 10). e) Cross-animal averaged response maps for the three response type groups (N = 10). Brain region contour lines are based on the ZBrain atlas: optic tectum (OT), torus longitudinalis (TL), habenula (Hab), cerebellum (Cer), tegmentum (Teg), oculomotor nucleus (nIII), trochlear nucleus (nIV), medial longitudinal fasciculus nucleus (nMLF), tangential nucleus (tVN), medial nucleus (mVN), inferior olive (IO), pretectum (PT), dorsal thalamus (DTh), ventral thalamus (VTh), and rhombomeres 1-7 (Rh 1-7). See also Figure S3, S4, S5 and S6.

Multilinear regression analysis identified three distinct neural response groups (Figure 3). The first group (blue) showed a mixed tonic-phasic response to nose-down pitch-tilts and a slight firing rate reduction for nose-up pitch-tilts. The second group (red) exhibited the opposite response, sensitive to nose-up tonic and phasic angular changes. The third group (yellow) displayed phasic responses to movements in both directions. Figure 3 presents these neurons’ locations in the representative example fish and a density plot summarizing the results obtained from N = 10 fish (individual results of all ten fish are shown in supplementary Figure S6). The nose-up responding neurons (orange) were mirror-symmetrically organized around the brain midline, predominantly in the tegmentum, with solid coverage of neurons of the nucleus of the medial longitudinal fasciculus (nMLF), the oculomotor nuclei (nIII), and the trochlear nucleus (nIV). They were also found in the pretectum, ventral and dorsal thalamus, lateral cerebellum, and rhombencephalon, including the inferior olive, tangential vestibular nucleus, and all rhombomeres. In contrast, the nose-down responsive neurons (blue) showed a similar mirror symmetric organization but with fewer neurons in the trochlear nucleus nIV and in the cerebellum and with a more lateral distribution in rhombomere 4-5. The distribution of the pure phasic response (yellow) encompasses neuronal populations especially in the torus semicircularis, optic tectum adjacent to the torus semicircularis, cerebellum, and rhombomeres 1–5. Importantly, this yellow cluster is spatially distinct and shows minimal overlap with the nose-up (red) and nose-down (blue) clusters. This indicates that the phasic response represents a separate population of neurons with a different anatomical and functional distribution compared to the direction-selective clusters. The cross-animal averaged response maps of the three response clusters exhibit a consistent spatial organization, indicating a highly stereotypic neuronal arrangement.

### Correlation Between Transgenic Line Expression and Vestibular Response Maps

By comparing our functional response maps with the transgenic line expression patterns collected in the Zebrafish Brain Browser [16], we have identified a significant overlap with the y249Et line’s expression profile. To delve further into this correspondence, we crossed this Gal4 line with a UAS line expressing the GCaMP7a sensor [17], revealing that a substantial number of the labeled neurons were responsive to vestibular stimulation (refer to Figure S1).

The y249Et line effectively labels neurons spanning a broad spectrum of brain regions characterized, where a substantial part of labeled neurons are responding to vestibular stimuli. Particularly noteworthy, marked and responsive neurons are distributed across the corpus cerebelli, eminentia granularis, nuclear medial longitudinal fascicle (nMLF), the oculomotor nuclei (nIII and nIV), the torus semicircularis, statoacoustic ganglion, vestibulospinal neurons, lateral reticular nucleus, inferior olive, and the interstitial nucleus of Cajal (a neuronal population rostral to the nMLF and not labeled in the zBrain atlas).

## Discussion

In this study, we conducted the first exhaustive brain-wide analysis of vestibular pitch-tilt response in larval zebrafish. We leveraged an advanced rotating light-sheet microscope for precise rotations along both the pitch-tilt and the roll-tilt axis. This approach enabled a detailed examination of the brain’s response to pitch-tilt stimulation and a comparative analysis of the roll-tilt response.

Our results reveal a widespread neuronal engagement throughout the brain during pitch-tilt stimulation, with neurons displaying a spatial arrangement that correlates with their sensitivity to specific tilt directions. This arrangement, mirror-symmetric across both brain hemispheres, suggests the presence of finely tuned neural circuits for distinct movement responses to pitch-tilt stimuli, offering insights into the sensorimotor processing involved. These findings, including a side-by-side comparison with roll-stimuli responses, are comprehensively illustrated in Figure 4.

**Figure 4:**
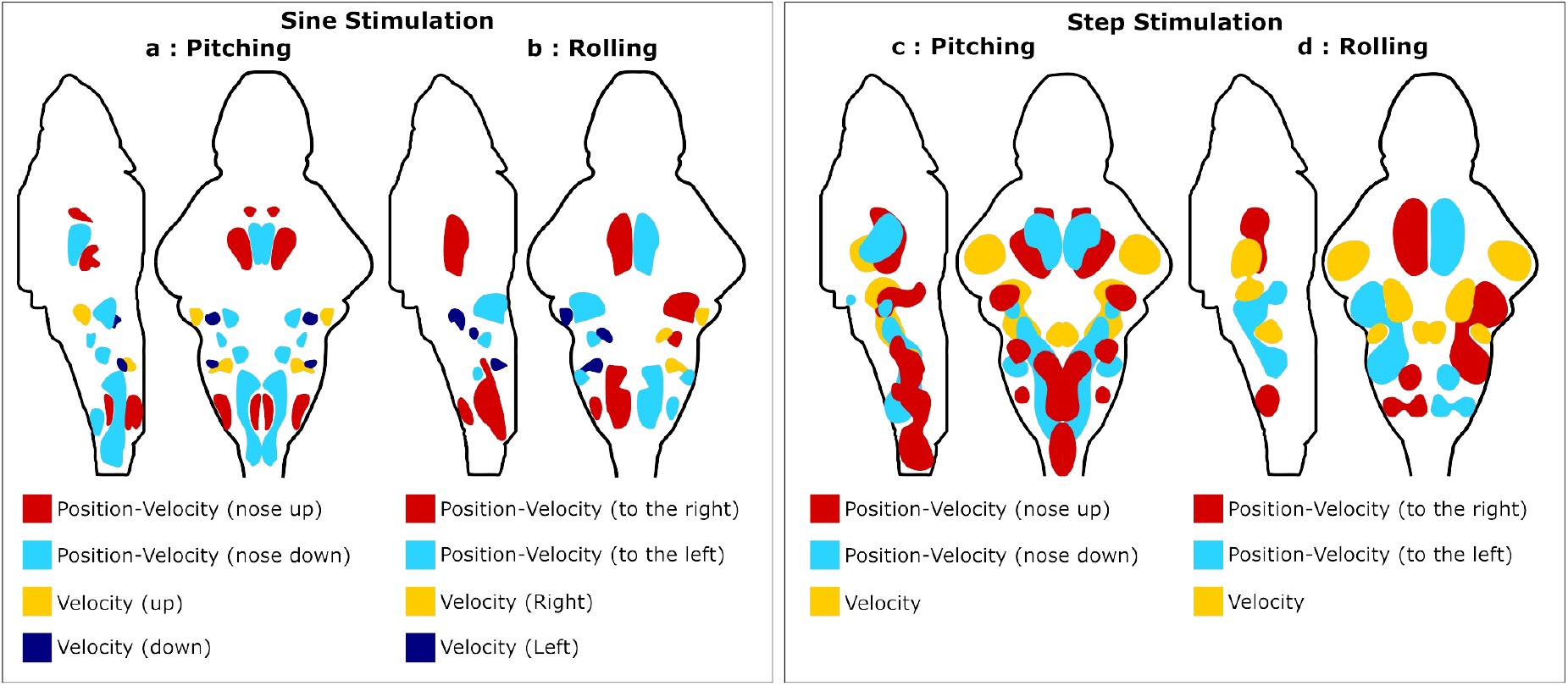
Summary of Brain Regions Responsive to Sinusoidal and Stepwise Stimulation along Pitch and Roll Axes. This graphical representation was established based on the following data: a) Figure 2e, right b) Figure 2f, right c) Figure 3e d) Figure 5 from Migault et al. 2018 [5].

### Directional Bias in Neuronal Recruitment During Pitch-Tilt Stimulation

#### Cerebellum

Our study highlights notable directional bias in brain region recruitment during pitch-tilt stimulation, particularly within the cerebellum. There was a pronounced cerebellar response to nose-up step stimuli, as opposed to a minimal average response to nose-down movements (Figure 3 e). This directional bias was apparent in both kernel density maps and neuron proportion histograms. By contrast, during the sinusoidal pitch-tilt stimulation, cerebellar neurons were only sparsely activated, indicating a response variance to the smoothness and acceleration of the stimulus.

The directional bias we observed in the cerebellar response, with more neurons responding to nose-up rotations, intriguingly mirrors the findings in vertical visual pursuit in primates and humans, where cerebellar Purkinje cells predominantly respond to downward visual motion [18, 19, 20]. Although one might expect minimal retinal image slip during an effective VOR, our recordings were performed in paralysed animals with the visual environment (the microscope) remaining static in the fish’s idiocentric reference frame. Thus, the observed directional bias likely reflects intrinsic vestibular processing. Under natural conditions, however, this intrinsic directional vestibular bias could interact with visual signals to contribute to gaze stabilization.

The inhibitory nature of Purkinje cells further suggests their role in facilitating downward eye movements by inhibiting muscles responsible for eye upward motion. This aligns with upward eye drifts seen in cerebellar atrophy patients [18] or primates with cerebellar lesions [21]. Furthermore, the consistent directional bias in cerebellar responses to both vestibular and visual stimuli points to multimodal convergence within the cerebellum. This aligns with the cerebellum’s known multimodal nature, highlighting its complex role in coordinating diverse sensory inputs. Furthermore, the observed cerebellar activity may also relate to the coordination of pectoral fin and trunk movements, which are known to contribute to lift generation during climbing behaviors in larval zebrafish [11, 22]. However, as our experimental setup did not include behavioral recordings of pectoral fin movements, we cannot directly address this aspect.

#### Oculomotor Neurons

Our study’s findings on the directionally biased activation of oculomotor neurons in nIII and nIV align with recent backtracing studies mapping these neurons to their specific muscle targets. [15]. The spatial organization of their responses reflects their role in controlling eye movements during pitch-tilt rotations. Notably, during nose-up rotations, nIV (innervating the superior oblique muscles) and the lateral part of nIII (affecting the inferior rectus muscles) coordinate for downward eye movement. In contrast, nose-down rotations involve the medial part of nIII, linked to the superior rectus and inferior oblique muscles, facilitating upward eye movements. These findings, illustrated in Figures Figure 2D and Figure S2, highlight the intricate neuromuscular control involved in ocular motion.

#### Caudal Hindbrain

In the caudal hindbrain, we also observed directionally biased recruitment of neurons: a medial-dorsal cluster responded to nose-up pitch-tilt movements, while two lateral strips were activated during nose-down movements. This differential activation pattern suggests the engagement of distinct postural motor programs, likely involved in compensating for destabilization in each direction. In particular, Ehrlich and Schoppik [22] have shown that pectoral fins generate lift during climbing trajectories but are not engaged during diving. Motor neurons controlling these fins are likely located in this region, making it plausible that the observed activation includes pectoral fin motor circuits. Simultaneous behavioral monitoring of pectoral fin engagement, or specific labeling of the corresponding motor neurons, could directly test this hypothesis. Together, these findings illustrate how the caudal hindbrain contributes to the biomechanical adjustments required for maintaining balance and posture along the pitch-tilt axis.

#### Vestibular Nucleus

Directionally biased neural recruitment within the vestibular nucleus has also been reported. Electrophysiological studies in cats revealed that 84% of secondary vestibular neurons are involved in vertical upward eye movements, in stark contrast to only 16% for downward movements [23]. In larval zebrafish, Schoppik et al. [24] mapped anatomical projections and reported a similar bias, with 85% of projection neurons targeting the oculomotor nucleus nIV, which drives downward movements. However, Goldblatt et al. [25] revisited this question using functional calcium imaging in the same preparation and reported a nearly equal distribution of nose-up and nose-down selective projection neurons (49% and 44%, respectively). Our functional mapping likewise reveals no strong directional asymmetry, aligning with the balanced distribution described by Goldblatt et al. Furthermore, Goldblatt et al. did not observe adirectional responses in the tangential nucleus under tonic or step stimuli. In our dataset, we counted only a small number of neurons with phasic, and thus potentially adirectional, responses. However, due to the absence of a specific genetic marker for tangential nucleus neurons in our experiments, our anatomical assignment relies on registration to the zBrain atlas and may therefore be less precise than targeted approaches. The small fraction of phasic neurons we identified may not be specific to the tangential nucleus.

Instead, we found differential activation of the medial and tangential vestibular nuclei in response to nose-down and nose-up step stimuli. This pattern is specific to step stimuli, as sinusoidal responses showed negligible activity in these nuclei. We cannot exclude that the elav3 line might predominantly label vestibular neurons with fast phasic responses, leading to their representation in step responses and potentially obscuring the projection asymmetry of secondary vestibular neurons in our functional maps. Future investigations employing transgenic lines that specifically label vestibular secondary neurons could provide deeper insights.

#### Vestibulospinal Neurons

Our study found vestibulospinal neurons on each side of the brain responsive to both pitch and roll stimuli, predominantly during nose-down tilts and ipsilateral ear-down rotations. However, it should be noted that these neurons might represent only a subset of all vestibulospinal neurons, as the elavl3 promoter does not label all neurons, and some vestibulospinal neurons may remain unlabeled. This directionally biased response aligns with anatomical studies demonstrating that these neurons predominantly receive input from hair cells in the macula, tuned to rostral-lateral otolith displacements [26]. Such responsiveness indicates their role in counteracting the postural challenges zebrafish typically encounter, particularly in situations of nose-down destabilization [12].

### Comparative Analysis of Pitch and Roll Tilt Responses

In our study, we conducted a comparative analysis of neuronal responses to pitch-tilt and roll-tilt stimuli, utilizing the same fish for both sets of stimuli for a sinusoidal stimulus waveform. Stepwise responses to roll-tilt stimuli have previously been characterized [5], and a qualitative comparison to our pitch-tilt step results confirms substantial differences in neuronal recruitment patterns across these two axes. The results of this comparison are visually depicted in Figure 4. This approach highlights the contrasting neuronal activation patterns in response to different types of vestibular stimulation, providing a unique perspective on how these stimuli are processed within the fish brain.

#### Hindbrain Oscillator

Our study revealed that neurons located medially in rhombomeres 1-3 are selectively activated by roll-tilt stimuli. Intriguingly, these neurons align with the hindbrain oscillator network [27, 28, 29] and evidence accumulation circuits [30, 31], previously attributed roles in integrating lateral visual flow for decision-making, especially in directional swimming during spontaneous exploration. This suggests that, alongside visual processing, these neurons also integrate vestibular information, indicating multimodal integration within these circuits. Such integration could have significant implications for precise postural control involving lateral tail movements. Notably, this activation pattern was absent in response to pitch-tilt stimuli, highlighting the response’s specificity.

#### Cerebellum

The cerebellum’s significant involvement during roll stimulation in fish underscores its essential role in maintaining a stable dorsal-up posture against destabilizing forces along the roll axis, particularly during swim bouts and turns. Unlike other movements, zebrafish larvae do not intentionally alter their body inclination along the roll axis for exploratory behavior. The cerebellum is likely continuously engaged in ensuring this constant dorsal-up orientation by modulating and refining motor actions. In contrast, body pitch angle control is less strict, often resulting in passive nose-down rotations of several degrees in pitch angle between consecutive bouts [12]. This might be reflected in the cerebellum’s relatively weaker recruitment during pitch-tilt vestibular stimulation and the low overlap between the two responses.

#### Phasic response

The observed distribution of neurons exhibiting pure phasic step responses provides new insight into the organization of vestibular processing circuits. We found that this response cluster includes populations within the torus semicircularis as well as several medial brainstem clusters that could not be precisely identified. Notably, the spatial distribution of these phasic-responsive neurons closely matches that observed for phasic roll responses in our previous work [5]. This overlap suggests that these neuronal populations are not only non-direction-selective with respect to pitch orientation (nose-up versus nose-down), but are also broadly responsive to different axes of rotation, including both pitch and roll. Such axis-independent, phasic encoding may reflect a shared or convergent pathway for rapid vestibular signaling across distinct postural movements. Furthermore, these neuronal populations show substantial overlap with those responsive to high-frequency auditory stimuli [32, 33]. This observation suggests that the phasic responses we detected are, at least in part, reactions to the onset of the step stimulus itself, which could induce vibrations or mechanical transients sensed primarily by the saccules. This may help explain the differential neural processing compared to the utricular low-frequency and tonic vestibular sensations captured by the other two step response clusters.

#### Overlap of Transgenic Line Expression Profiles with Vestibular Functional Maps

In our study, we also explored the correlation between our vestibular functional response maps and the expression patterns of various transgenic fish lines collected in the Zebrafish Brain Browser ([16]). This led to the identification of the yt249 transgenic line showing significant overlap with our functional maps. This line holds potential for future studies in tracing, optogenetics, or targeted ablations, which are crucial for a deeper understanding of the neural circuits involved in vestibular processing.

Our study advances the understanding of differential neuronal responses and asymmetries in pitch and roll-axis vestibular stimulation. Through extensive brain-wide assessments in larval zebrafish and analysis of transgenic fish line overlaps, we contribute to the growing knowledge of the vestibular system. The insights from this research are significant for comprehending postural control, spatial orientation, and the broader implications for neurological disorders associated with vestibular dysfunction [34].

## METHODS

### Animal Husbandry

All experiments were performed on 6-7 dpf larvae. The sex of the animals was not yet determined at this age, and was therefore not reported. Adult fish were maintained at 28°C in system water (pH 7-7.5 and conductivity between 300 and 350 *µ*S) in the fish facility of the Institut de Biologie Paris-Seine. Eggs were collected in the morning and then kept in a Petri dish with E3 at 28°C under a 14h/10h light/dark cycle. Larvae were kept in groups of 10-40 fish and fed powdered food from 5 dpf on. Calcium imaging experiments were carried out in two different transgenic lines: *elavl3:H2B-GCaMP6f* [35] and *y249:Gal4; UAS:GCaMP7a* [16, 17], both in nacre background. We have complied with all relevant ethical regulations for animal use. Experiments were approved by Le Comité d’Éthique pour l’Expérimentation Animale Charles Darwin C2EA-05 (02601.01 and #32423-202107121527185 v3).

### 3D Printing

The bracket and all other custom parts (1B, cyan elements) were printed using an Original Prusa MINI+ 3D printer, and we used Prusament PETG filament with a layer quality of 0.2mm and a density of 80%. The 3D design file is open source and can be accessed at https://a360.co/2U1KBoH.

### Sample Preparation

First, the fish were paralyzed by immersing them for 75 seconds in 25 *µ*L of a 1 mg/mL *α*-bungarotoxin solution (Thermo Fisher Scientific) in E3 medium. They were then transferred to pure E3 medium and left for 30 minutes to ensure the absence of motor activity while maintaining a regular heartbeat. Subsequently, the larvae were carefully placed in a 2% low-melting agarose solution (Sigma-Aldrich) and tail-first drawn into a 1 mm inner diameter glass capillary tube using a plunger. After this step, the agarose was cut directly rostral to the fish.

Fish were mounted as close to horizontal as possible along the pitch axis, with the average orientation estimated at −3.9^°^ *±* 4.0^°^ (mean *±* SD; nose-down). This orientation is well within the natural range of pitch postures observed in freely swimming larvae (9^°^ *±* 13^°^, nose-up; [12]) and is small compared to the *±*20^°^ stimulation range used in our experiments. Although freely swimming larvae on average exhibit a slightly nose-up posture, the variability is substantial. We therefore used the horizontal orientation as our experimental reference to provide a reproducible and consistent baseline.

### Acquisition Protocol

We conducted volumetric imaging at a rate of 2.5 stacks per second, covering 25 brain sections. Each section had an inter-slice distance of 10 *µ*m and a pixel size of 1.2×1.2 *µ*m^2^.

### Quantification and Statistical Analysis

For the characterization of the setup mechanical stability during microscope rotation, the processing of functional light-sheet data, the registration onto the z-Brain atlas, the generation of the phase maps, the regression analysis, and the generation of the average cluster density maps, we followed the protocols described in Migault et al. [5]. Any modifications to these procedures are detailed in the corresponding sections below. Data analysis was performed in MATLAB. The runquantile function from the caTools R package (https://CRAN.R-project.org/package=caTools) was accessed from within MATLAB. Morphometric registration was conducted using the Computational Morphometry Toolkit (CMTK).

### Phase Maps

First, we estimated the exact stimulation frequency and its phase from the Fourier spectrum of the motor monitor signal. Then, we calculated the power spectrum of the fluorescent time trace for every voxel in the recorded 4D brain stack. We estimated amplitude and phase at the stimulation frequency. We normalized the amplitude to a signal-to-noise value by subtracting the spectrum baseline from the estimated peak height at the stimulation frequency and dividing this difference by the baseline value. The baseline was estimated as the mean value over a window to the right of the peak. Finally, we estimated the phase shift of the fluorescent response relative to the stimulus by subtracting the motor monitor phase from the fluorescent signal phase.

### Decay Time Constant of Calcium Sensor

In our previous work, we estimated the phase shift induced by the GCaMP6s reporter to be −1.35 *±* 0.05 rad [5]. In this study, we estimated the phase shift for the GCaMP6f reporter by comparing neural response phases in the trochlear nucleus during rolling stimulation with those from larvae expressing GCaMP6s. Due to its consistent and significant neuronal responses, the trochlear nucleus was chosen for this comparison. The observed phase shift difference between GCaMP6s and GCaMP6f was 0.20 *±* 0.05 radians. Consequently, we deduced the phase shift introduced by GCaMP6f to be −1.15 *±* 0.10 radians. This translates to a decay time of *τ* = −*ω*^−1^ tan *ϕ* = 1.78 *±* 0.80 seconds, significantly smaller than the decay time of 3.5 *±* 0.7 seconds for GCaMP6s, as we previously estimated.

### Assessment of Linear Acceleration Due to Off-Axis Mounting

To rotate the microscope, we use a high-precision, high-load motor with PID-controlled movement. The maximal angular acceleration was set to 1500 ^°^*/*s^2^. For sinusoidal rotation, the angular acceleration is sinusoidal and in phase with the stimulus. The peak angular acceleration is

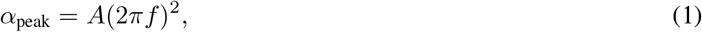

with *A* the rotation amplitude (in radians) and *f* the frequency. For our waveforms (*A* = 20^°^ = 0.349 rad, *f* = 0.2 Hz), this yields

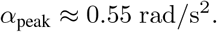

The fish is mounted as close as possible to the rotation axis, with a residual off-center distance *r <* 1 mm. Such off-center positioning produces a tangential linear acceleration

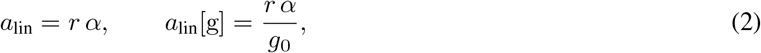

where *g*_0_ = 9.81 m*/*s^2^. Under our sinusoidal conditions and *r* =1 mm,

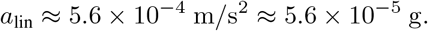

Even at the motor’s maximal acceleration 1500 ^°^*/*s^2^ (*α* = 26.18 rad*/*s^2^), the linear acceleration remains modest:

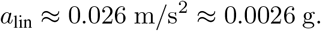

This peak occurs only transiently during the step protocol as the microscope accelerates to the ramp velocity of 30 ^°^*/*s. For intuition, a lateral linear acceleration *a*_lin_ is otolith-equivalent to a static roll tilt *θ* given by

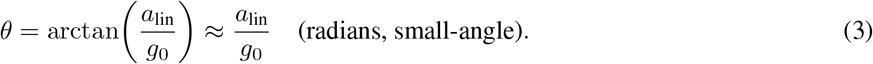

Thus, our maximal *a*_lin_ ≈ 0.0026 g corresponds to *θ* ≈ 0.0026 rad ≈ 0.15^°^. From this analysis, linear acceleration due to off-axis mounting can be neglected; the vestibular response is dominated by gravitational acceleration.

### Quantification of misalignment-induced cross-talk

We used the following convention to calculate the crosstalk resulting from mounting misalignment between the fish and the rotation stage.

#### Reference frames

The *world (stage) frame* (*x*_*w*_, *y*_*w*_, *z*_*w*_) is defined with *z*_*w*_ pointing upwards. The *fish frame* (*x*_*f*_, *y*_*f*_, *z*_*f*_), initially aligned with the world frame, is defined such that *x*_*f*_ points left–right (toward the fish’s left), *y*_*f*_ points caudal–rostral (with rostral being +*y*_*f*_), and *z*_*f*_ points ventral–dorsal (with dorsal being +*z*_*f*_).

#### Mounting offsets

Misalignment was parameterized as a roll offset (rotation about world *y*_*w*_) by angle *ϕ*, pitch offset (rotation about word *x*_*w*_) by angle angle *γ*, and a yaw offset (rotation about world *z*_*w*_) by angle *θ*, applied sequentially as

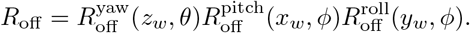

The corresponding rotation matrices are

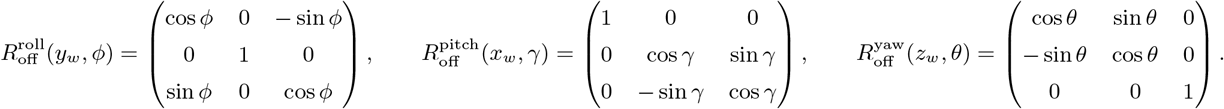

#### Stage commands

A *pitch command* corresponds to a rotation about world *x*_*w*_ by angle *α*, and a *roll command* to a rotation about world *y*_*w*_ by the same angle *α*, through

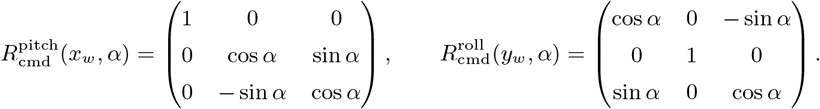

#### Projection of gravity

The gravitational acceleration in the world frame is **g**_*w*_ = (0, 0, −*g*). We projected this vector into the fish reference frame to estimate the gravitational force components experienced by the otoliths during commanded stage rotations as a function of the misalignment angles:

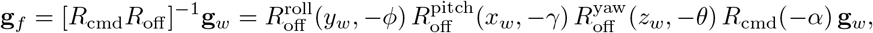

where *R*_cmd_ denotes either a commanded pitch or roll rotation.

#### Resulting gravity components

Evaluating these expressions gives

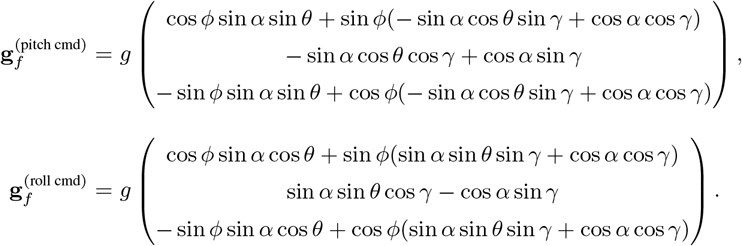

#### Definition of crosstalk fraction

We defined the crosstalk fraction as the relative magnitude of the unintended gravitational component (along the orthogonal axis) to the intended one:

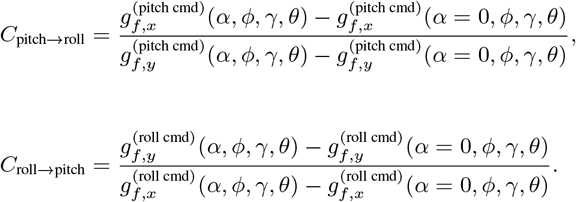

#### Crosstalk estimate

Table 1 shows the crosstalk fractions for all fish, estimated for the maximal commanded rotation amplitude (*α* = 20^°^). Residual cross-axis contamination remained small (with *C*_pitch→roll_ = 3.98 *±* 3.66 % (N=10, mean, std) and *C*_roll→pitch_ = −3.19 *±* 3.74 % (N=8, mean, std)). Thus, the partial overlap of neuronal responses to roll- and pitch-axis stimulation observed in the main text cannot be attributed to mechanical or geometric cross-talk, but reflects genuine axis-independent vestibular sensitivity.

**Table 1:** Misalignment angles and resulting crosstalk in percent shown for all fish. Fish one is the single fish example displayed in Figure 2 and 3.

| Fish | Roll config |  |  |  | Pitch config |  |  |  |
| --- | --- | --- | --- | --- | --- | --- | --- | --- |
|  | Misalignment angle (deg) |  |  | Crosstalk (%) | Misalignment angle (deg) |  |  | Crosstalk (%) |
| | $\varphi$ | $\gamma$ | $\theta$ | | $\varphi$ | $\gamma$ | $\theta$ | |
| 0 | 3.58 | 0.74 | -4.10 | -7.03 | 2.76 | 0.38 | -3.76 | 7.45 |
| 1 | 1.83 | 0.07 | -0.09 | -0.13 | -1.33 | 0.11 | 0.92 | -1.19 |
| 2 | -0.33 | 0.23 | -4.77 | -8.26 | -0.12 | 0.07 | -5.37 | 9.36 |
| 3 | -2.44 | 0.05 | -2.93 | -5.06 | 0.85 | 0.17 | -3.13 | 5.74 |
| 4 | -2.96 | 0.37 | -3.43 | -5.94 | 1.97 | 0.15 | -3.00 | 5.86 |
| 5 | -2.98 | 0.13 | 0.06 | 0.15 | 3.90 | 0.03 | 0.98 | -0.51 |
| 6 | 1.27 | 0.14 | -0.42 | 0.79 | 2.71 | -0.00 | 1.26 | -1.36 |
| 7 | -1.95 | 0.37 | -0.09 | -0.05 | 2.37 | 0.66 | -2.59 | 5.29 |
| 8 |  |  |  |  | 2.18 | 0.21 | -0.75 | 1.99 |
| 9 |  |  |  |  | 1.90 | 0.50 | -2.87 | 5.62 |
| 10 |  |  |  |  | -1.84 | 0.02 | -3.50 | 5.56 |

### Regression Analysis

We conducted the multilinear regression analysis using a modified version of the protocol from Migault et al. (2018). In addition to two angular position regressors, we introduced four new regressors based on the microscope’s velocity signal. These regressors, defined by the angular velocity’s direction, differentiate between movements away from or towards the preferred dorsal-up orientation (see Figure S3). This enhanced the analysis’s precision. Neurons were classified as responsive if their regression coefficients and T-scores were above the 94th percentile for at least one of the six regressors. The analysis involved convolving all six regressors with the H2B-GCaMP6f response kernel, estimated as a single exponential decay with a *τ* = 1.78 s decay time (see Figure S3). To identify highly responsive neurons, we set a threshold at the 94th percentile for both the regression coefficient and T-score distributions. Neurons within the intersection of both 94th percentiles were clustered, indicating a robust response to the regressor (manifested as a high coefficient) and a significant explanation of neuron variance (reflected in high T-scores). This selection procedure is illustrated in supplementary Figures S4 and S5 illustrate this selection procedure.

### Statistics and reproducibility

Sample sizes were determined by the number of successful recordings obtained and the extent of completed downstream analysis, rather than by a formal a priori power calculation. Data were analyzed in chronological order of acquisition. In total, 12 fish were recorded under sinusoidal roll-tilt and pitch-tilt stimulation, of which the first 8 chronological recordings were processed through the full analysis pipeline and included in the sinusoidal population analyses reported here. Datasets not included in the reported analyses correspond to the most recent recordings and were omitted because of the substantial analysis burden, rather than being excluded on the basis of response quality or outcome. For these sinusoidal experiments, no separate control group was used because the main comparison was between stimulation conditions tested in the same animals (within-subject/repeated-measures design). Stepwise pitch-tilt responses were recorded in all but the first fish, yielding 11 fish in total; of these, the first 10 chronological recordings were processed through the full analysis pipeline and included in the stepwise population analyses reported here. The dataset not included in the reported analysis corresponded to the most recent recording and was omitted because of the substantial analysis burden, rather than being excluded on the basis of response quality or outcome. In the present study, only pitch step responses were quantified. Roll step responses were not recorded in the same animals for direct comparison; previously published roll step-response data obtained in a separate set of fish were used for qualitative comparison.

Fourier-based phase maps were computed per pixel via Fourier analysis of 150-cycle sinusoidal trials for each larva (N = 8; fish per batch: 3, 2, 3), then morphometrically registered to the Z-Brain reference (CMTK) and averaged pixel-wise across larvae. For step stimuli, ordinary least squares regression was fitted per neuron in each larva (N = 10; fish per batch: 2, 2, 5, 1), yielding coefficients and T-scores (*β* ≠ 0, [36]). No formal normality tests were performed on residuals; with *>* 1000 time-points per neuron, coefficient estimates approximate normality by the central limit theorem. P-values and degrees of freedom were not calculated; T-scores alone were used for neuron selection. Highly responsive neurons were defined by a one-sided cutoff at the 95th percentile of both coefficient and T-score distributions (see figure in supplementary). Cluster densities were estimated per fish via Gaussian KDE (bandwidth = 12.8 *µ*m) on a 5.5 *µ*m isotropic grid, and final group-level maps were obtained by elementwise averaging these per-fish KDE maps, thereby preserving fish-level independence. Stimulus direction and velocity were included as covariates; larval sex and age (6–7 dpf) were not tested. All summary statistics are reported as mean ± SD; formal 95 % confidence intervals, Bayesian analyses, and effect-size metrics (e.g. Pearson’s r, R^2^) were not calculated.

## Resource Table

**Table 2:** Key resource table.

| REAGENT OR RESOURCES | SOURCE | IDENTIFIER |
| --- | --- | --- |
| **Chemicals** |  |  |
| 2D Galvo Mirror Systems | Thorlabs | GVS01@ |
| Low melting point agarose | Sigma-Aldrich | A9414-50G |
| α-bungarotoxin | ThermoFisher Scientific | B-1601 |
| **Experimental Models: Organisms/Strains** |  |  |
| Tg(elav3-H2B:GCaMP6f) |  | Vladimirov et al., 2014 [35] |
| y249:Gal4 ; UAS:GCaMP7a |  | Marquart et al., 2015 and Muto et al., 2013 [16, 17] |
| **Software and Algorithms** |  |  |
| Matlab | The MathWorks | https://www.mathworks.com/products.html |
| Labview | National Instruments | https://www.ni.com |
| CMTK | Rohlfing and Maurer, 2003 [37] | https://www.nitrc.org/projects/cmtk/ |
| FIJI | (ImageJ) NIH | https://fiji.sc/ |
| Z-Brain atlas | Randlett et al., 2015 [38] | https://zebrafishexplorer.zib.de/home/ |

## RESOURCE AVAILABILITY

### Lead contact

Further information and requests for resources and reagents should be directed to and will be fulfilled by the lead contact, Volker Bormuth.

### Data availability

Raw data will be shared by the lead contact upon request. Numerical source data for graphs and charts are available at 10.5281/zenodo.19253364 [39].

The key resource table lists all the fish lines used in this study. This study did not produce new transgenic lines; all the lines used have already been published and are available upon request. This study did not generate further new unique reagents.

### Code availability

Code will be shared by the lead contact upon request.

### Author Contributions

G.M. and V.B. conceived the project, G.M. and V.B. built the instrument, G.M., N.B., S.C., V.B. performed experiments, G.M., N.B., S.C., G.D., V.B. analyzed data, G.M., N.B., S.C., G.D., V.B. wrote the manuscript.

## ACKNOWLEDGEMENTS

We thank the IBPS fish facility staff for their invaluable help. We are also grateful to Carounagarane Dore for his important contribution to the realization of the mechanical parts of the light-sheet microscope. We thank Misha Ahrens for providing the GCaMP6f line, Koichi Kawakami for the UAS:GCaMP7a line, and Harold Burgess for the yt249 line.

## Funding Statements

This project has received funding from the European Research Council (ERC) under the European Union’s Horizon 2020 research and innovation grant agreement number 715980 and by the ATIP-Avenir program from CNRS and INSERM. G.M. was funded by a PhD fellowship from the Doctoral School in Physics, Ile de France (EDPIF). S.C. received funding from the European Union’s Horizon 2020 research and innovation program under the Marie Skłodowska-Curie grant agreement number 813457 ZEbrafish Neuroscience International Training Hub (ZENITH) as well as the Fin de thèse grant from the Fondation pour la Récherche Medicale (FRM). The funders had no role in study design, data collection, analysis, interpretation, manuscript preparation, or the decision to submit the work for publication.

## DECLARATION OF INTERESTS

The authors declare no competing interests.

## 1 Videos legends

**Video S1 Trial-averaged response to sinusoidal pitch-tilt stimulation**

**Video S2 3D animated average phase map to pith-tilt stimulus**.

**Video S3 3D animated average phase map to roll-tilt stimulus**.

## 2 SUPPLEMENTARY FIGURES

### Correlation Between Transgenic Line Expression and Vestibular Response Maps

Compared to the phase maps shown in Figure 2, the calcium sensor used here is GCaMP7 rather than GCaMP6f, and it is expressed in the cytoplasm instead of being restricted to the nucleus. These differences affect both the sensor’s kinetics and the dynamics of intracellular calcium, potentially resulting in phase offsets. Additionally, the cytoplasmic expression in the y249Et line enables visualization of axonal projections, which are not labeled in nuclear-targeted lines.

**Figure S1:**
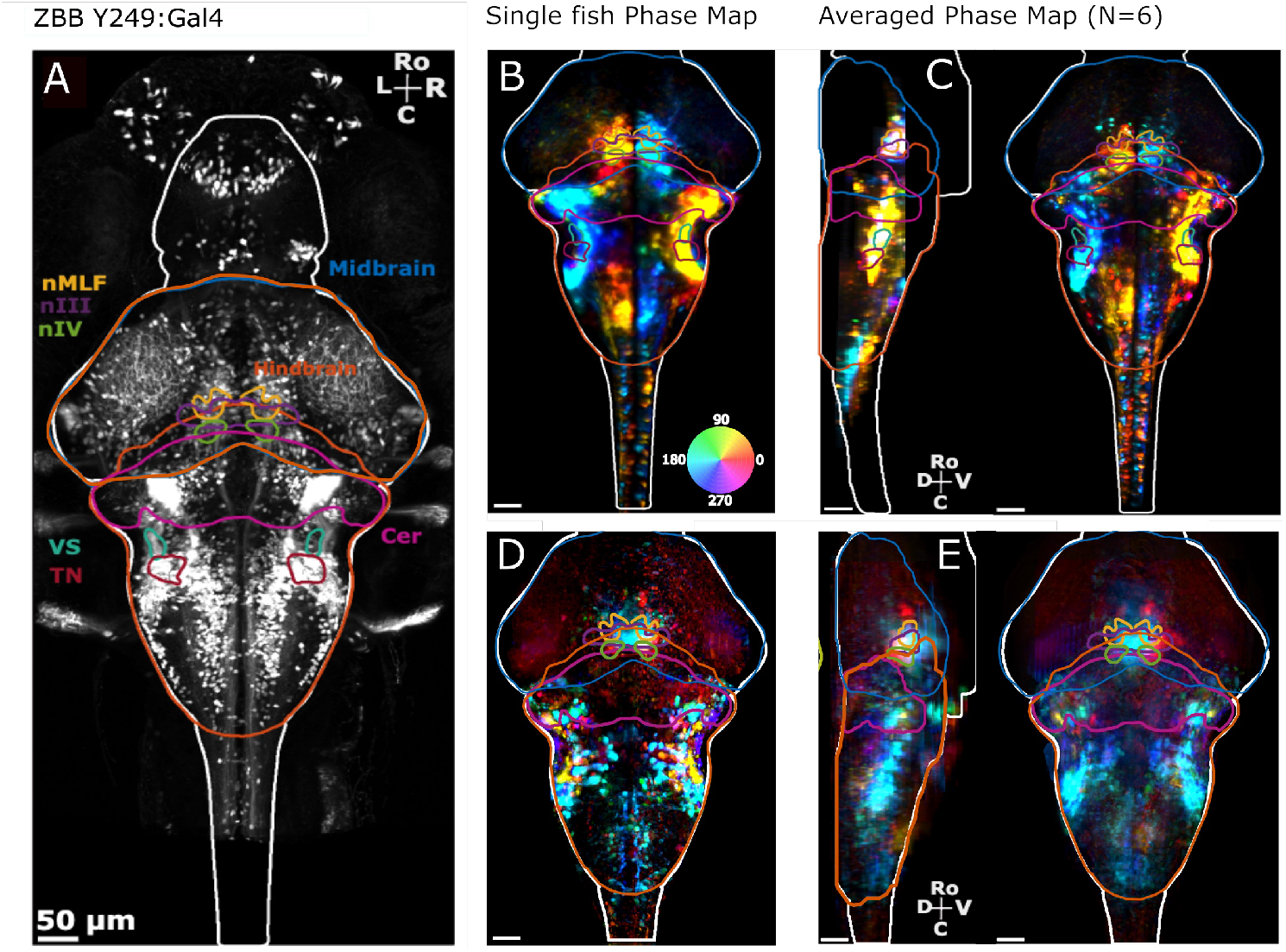
Functional Brain-wide Response to Sinusoidal Roll in the y249Et Line. (A) Z-projection of the anatomical stack of the Y249Et transgenic line downloaded from the ZebraBrainBrowser library and mapped onto the zBrain atlas. (B) Z-projection example-fish phase map displaying neurons active during sinusoidal roll. Yellow-orange regions indicate activity during right-ear-down roll, while blue and cyan indicate activity during left-ear-down roll. (C) Side and dorsal views of the average phase map (N = 6 fish). (D) Z-projection example-fish phase map displaying neurons active during sinusoidal pitch-tilt stimulation. Red regions indicate activity during nose-up pitch, while cyan indicates activity during nose-down pitch. (E) Side and dorsal projections of the average phase map (N = 10 fish). Brain region contour lines are based on the ZBrain atlas: cerebellum (Cer), oculomotor nucleus III (nIII), trochlear nucleus IV (nIV), medial longitudinal fasciculus nucleus (nMLF), tangential nucleus (TN), and vestibulospinal population (VS). Scale bar = 50 *µ*m.

**Figure S2:**
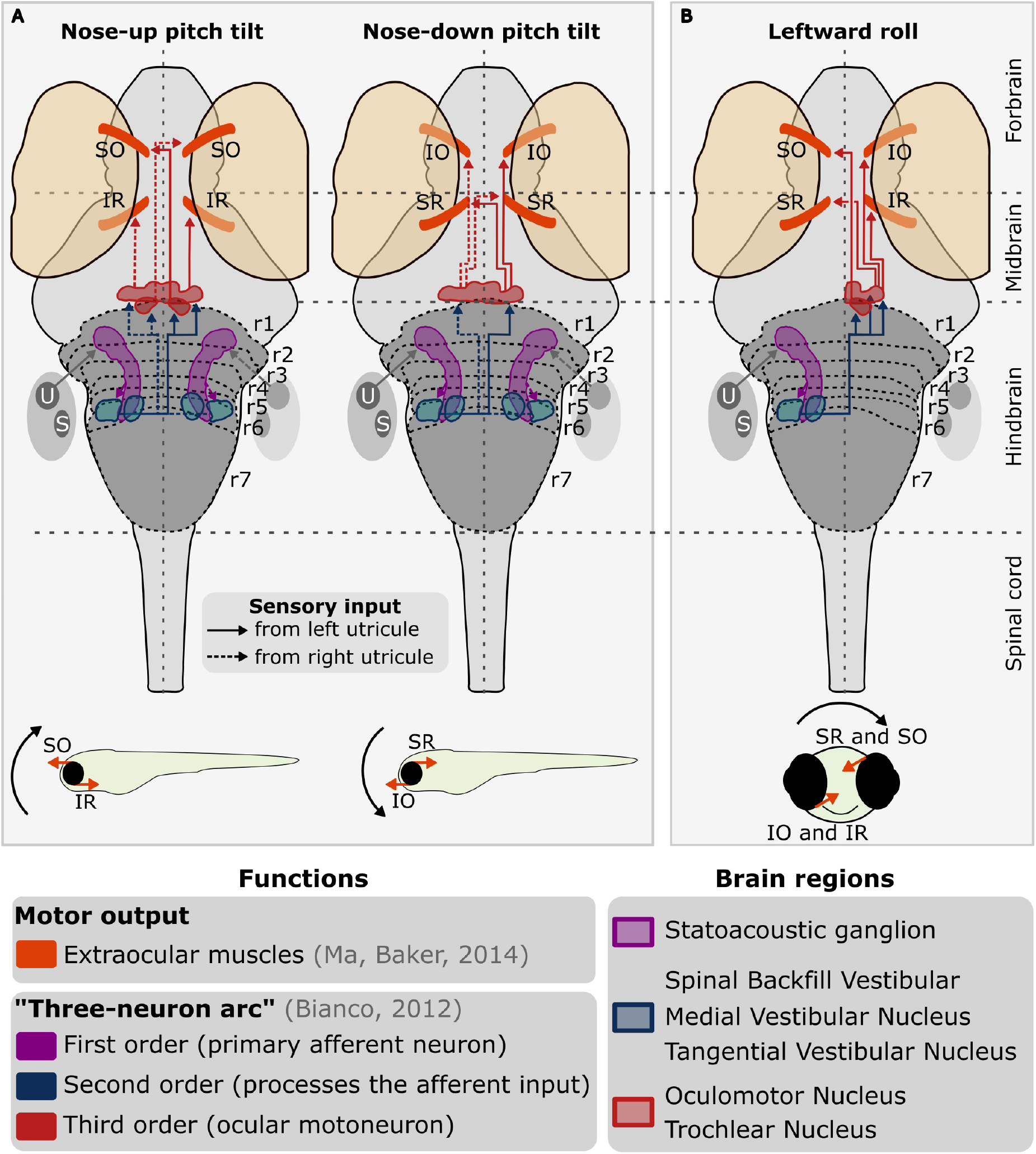
Schematic of Neural Network and Behavioral Responses during Vestibulo-Ocular Reflex (VOR). Related to Figure 2. Schematic representation of the neural network involved in the angular vestibulo-ocular reflex (VOR) during pitch-tilt (A) and roll-tilt (B), accompanied by corresponding behavioral response patterns (black arrow indicates stimulation direction, while orange arrows depict the force direction exerted by the extraocular muscles). Key elements include the extraocular muscles (superior rectus, SR; inferior rectus, IR; inferior oblique, IO; and superior oblique, SO), utricle, U; saccule, S; and rhombomeres, r1 to r7.

**Figure S3:**
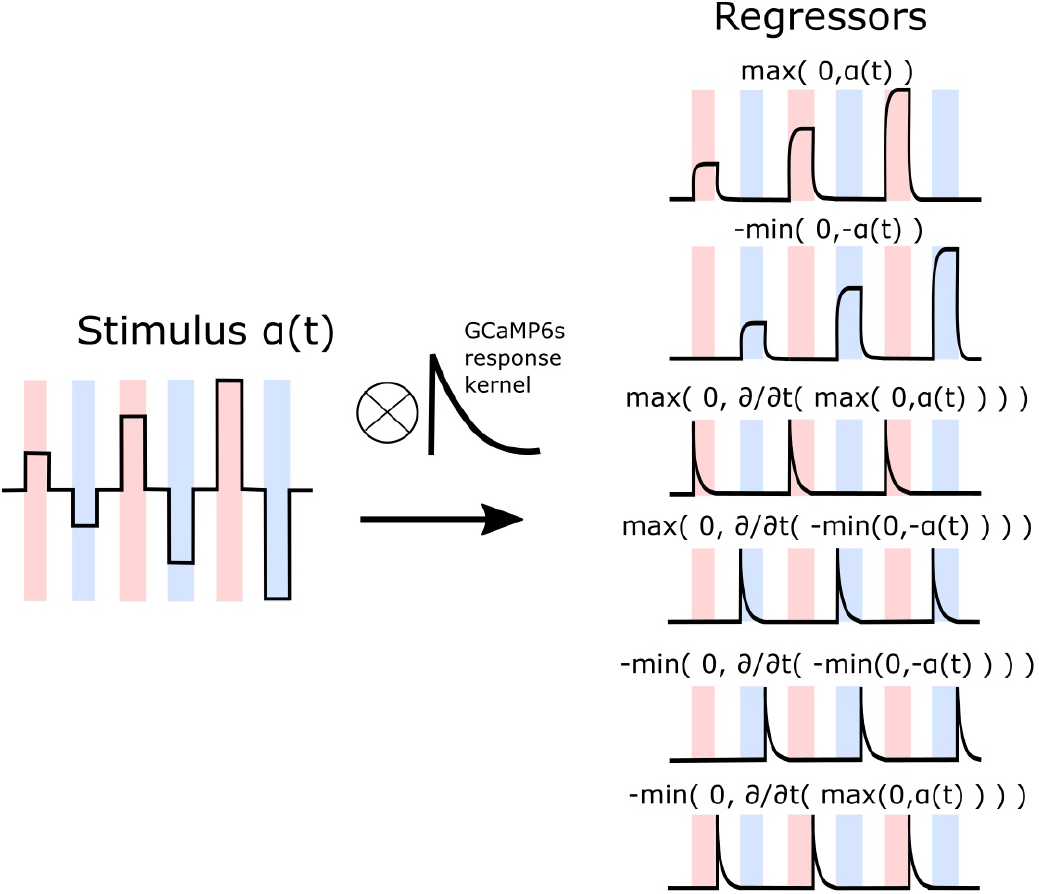
Construction of Regressors from the Stimulus. Related to Figure 3. This graphical representation demonstrates the sequential process of convolving the stimulus signal, *ω*(*t*), with the kernel representing the temporal dynamics of the GCaMP calcium sensor. The subsequent creation of positive tonic and phasic components is depicted, resulting in the formation of the six distinct regressors utilized in the multilinear regression analysis.

**Figure S4:**
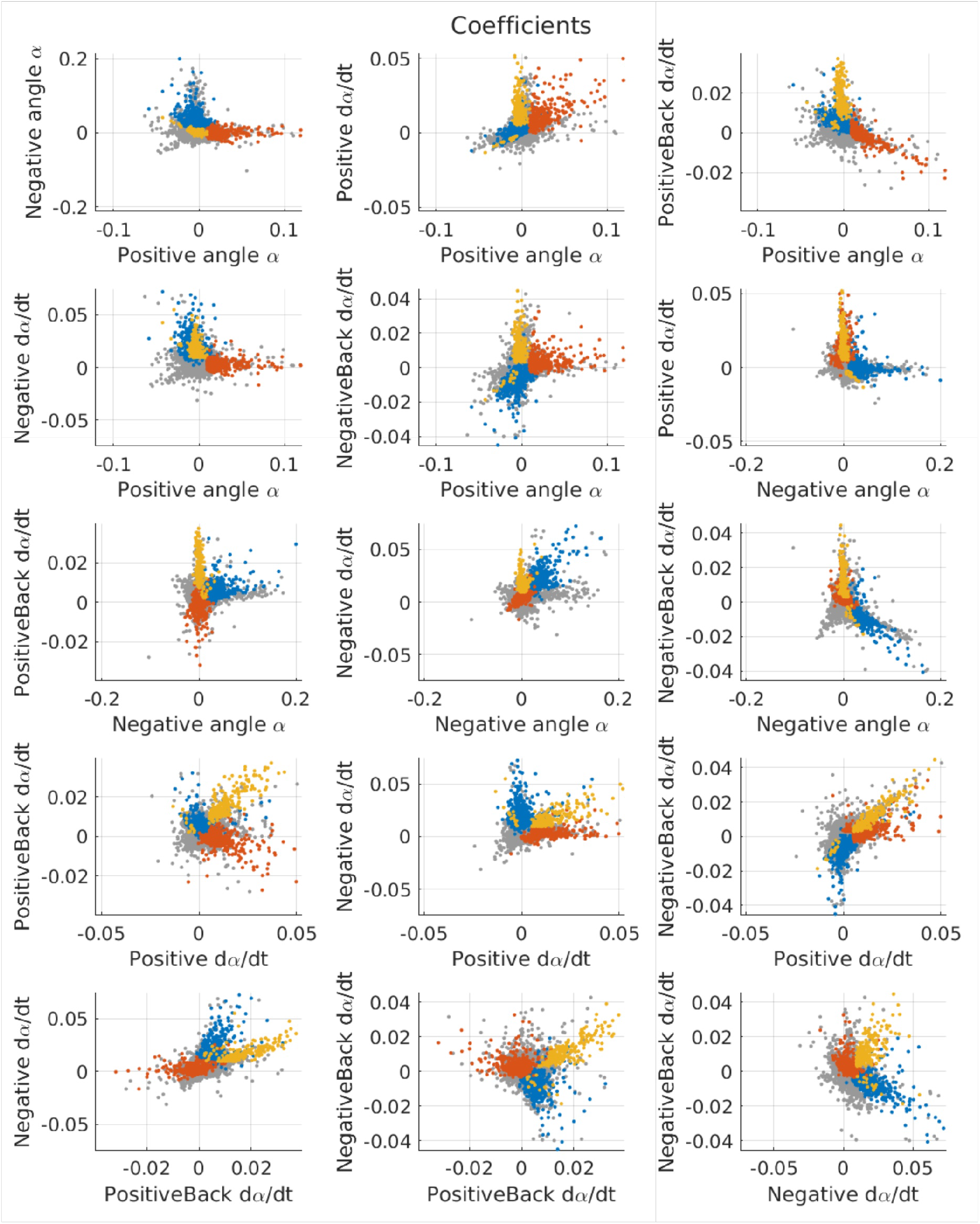
Scatter Plots of Regression Coefficients for Identified Neuronal Response Groups. Related to Figure 3. Scatter plots representing the regression coefficients corresponding to the three identified groups of neuronal responses, as illustrated in Figure 3. Gray dots denote neurons not categorized within these three groups.

**Figure S5:**
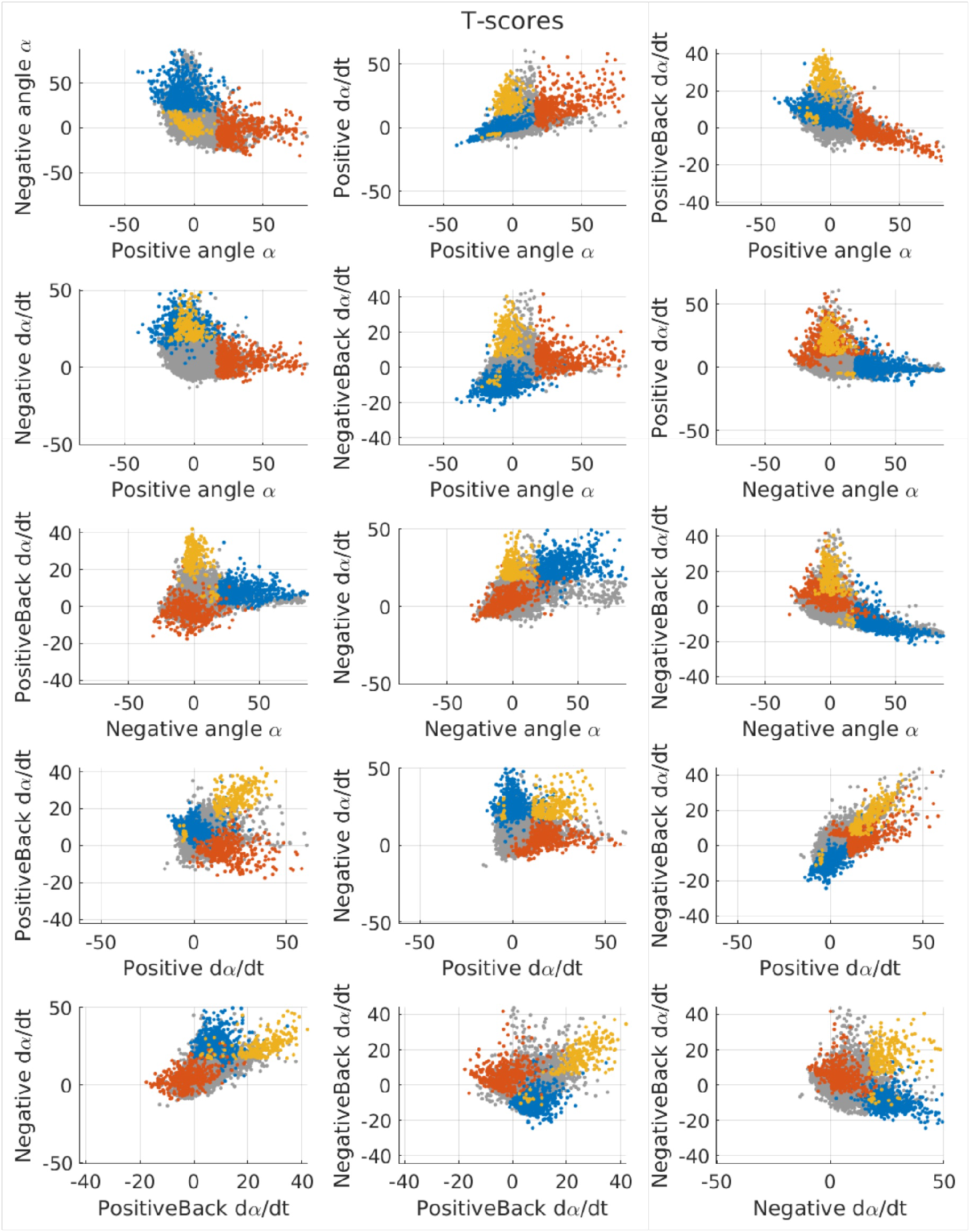
Scatter Plots of Regression T-Scores for Identified Neuronal Response Groups. Related to Figure 3. Scatter plots depicting the regression T-scores associated with the three identified groups of neuronal responses, as depicted in Figure 3. Neurons falling outside of these three groups are represented as gray dots.

**Figure S6:**
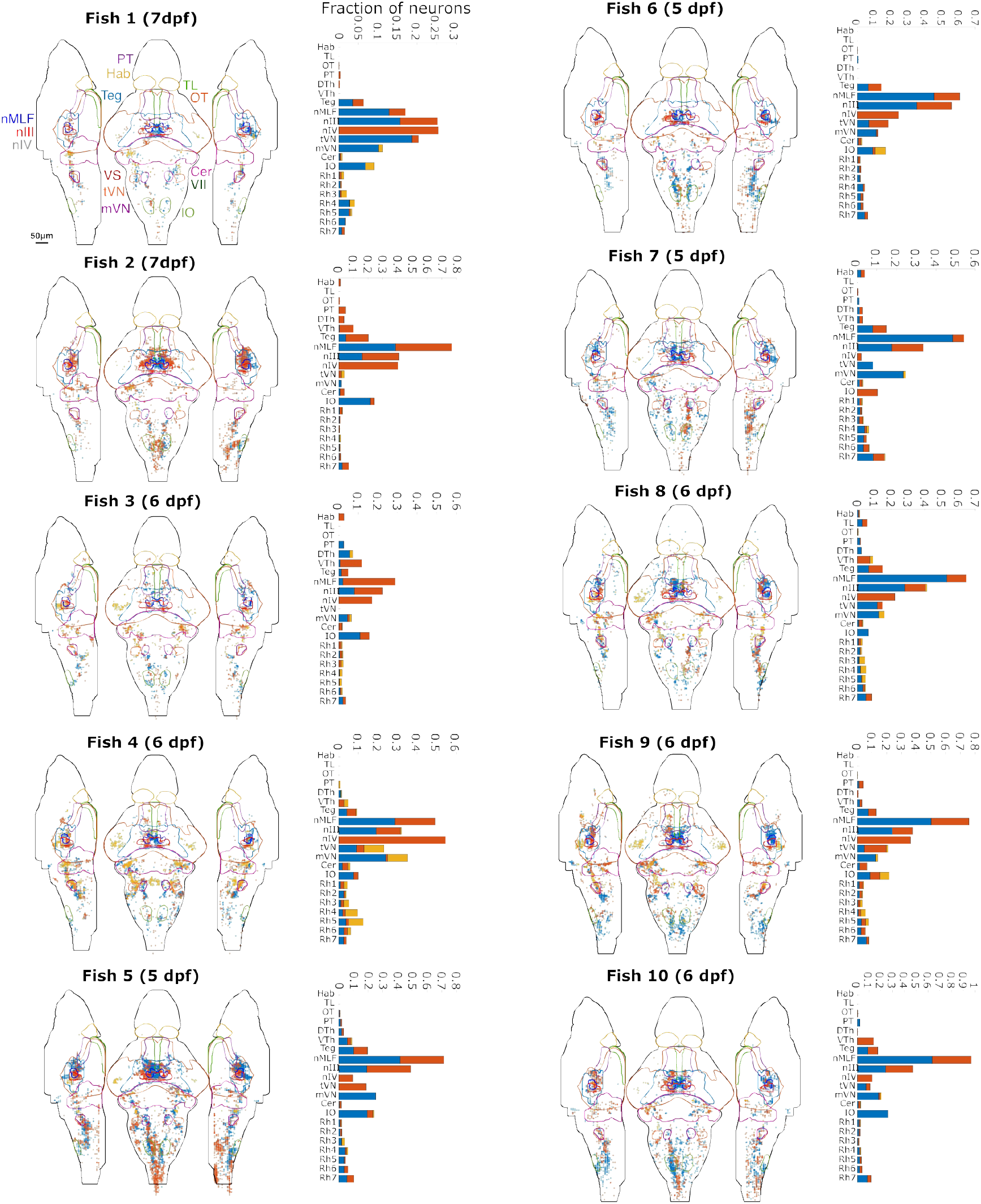
Regression Results Visualization and Histograms of Neuronal Response Groups. Related to Figure 3. Visualizations of the regression results for the ten fish used in the average are depicted in Figure 3. Accompanying histograms illustrate the fraction of neurons belonging to the three identified groups within specific brain regions.

## Notes

### Competing Interest Statement

The authors have declared no competing interest.

### Summary of Updates

This revised manuscript incorporates major changes to improve methodological clarity and interpretation. We added new methodological controls, including an estimate of linear acceleration due to off-axis mounting and a quantitative analysis of misalignment-induced pitch-roll crosstalk. We also revised the quantitative analysis by excluding vestibulospinal neurons from the step-response regional histograms because anatomical assignment in this small region was not sufficiently robust for quantitative comparison. In addition, the Results and Discussion were substantially reorganized and refined to better distinguish direction-selective and phasic response populations, strengthen the comparison between pitch and roll processing, and correct the discussion of oculomotor responses while aligning the interpretation more carefully with prior literature. We further clarified and expanded key aspects of the experimental design, anatomical assignments, figures, and legends. Finally, the open-source microscope design files were deposited in Zenodo for persistent public access.

## References

1. Dale, M. L., Horak, F. B., Wright, W. G., Schoneburg, B. M., Nutt, J. G., and Mancini, M. (2017). Impaired perception of surface tilt in progressive supranuclear palsy. PLoS ONE 12, e0173351. 10.1371/journal.pone.0173351.

2. Cronin, T., Arshad, Q., and Seemungal, B. M. (2017). Vestibular Deficits in Neurodegenerative Disorders: Balance, Dizziness, and Spatial Disorientation. Frontiers in Neurology 8, 538. 10.3389/fneur.2017.00538.

3. Brandt, T., Strupp, M., and Dieterich, M. (2014). Towards a concept of disorders of “higher vestibular function”. Frontiers in Integrative Neuroscience 8, 47. 10.3389/fnint.2014.00047.

4. Mansour, Y., Burchell, A., and Kulesza, R. J. (2021). Central Auditory and Vestibular Dysfunction Are Key Features of Autism Spectrum Disorder. Frontiers in Integrative Neuroscience 15, 743561. 10.3389/fnint.2021.743561.

5. Migault, G., Plas, T. L. v. d., Trentesaux, H., Panier, T., Candelier, R., Proville, R., Englitz, B., Debrégeas, G., and Bormuth, V. (2018). Whole-Brain Calcium Imaging during Physiological Vestibular Stimulation in Larval Zebrafish. Current Biology 28, 3723–3735.e6. 10.1016/j.cub.2018.10.017.

6. Hamling, K. R., Zhu, Y., Auer, F., and Schoppik, D. (2023). Tilt in Place Microscopy: a Simple, Low-Cost Solution to Image Neural Responses to Body Rotations. The Journal of Neuroscience 43, 936–948. 10.1523/jneurosci.1736-22.2022.

7. Tanimoto, M., Watakabe, I., and Higashijima, S.-I. (2022). Tiltable objective microscope visualizes selectivity for head motion direction and dynamics in zebrafish vestibular system. Nature communications 13, 7622. 10.1038/s41467-022-35190-9.

8. Sugioka, T., Tanimoto, M., and Higashijima, S.-I. (2022). Biomechanics and neural circuits for vestibular-induced fine postural control in larval zebrafish. Nature communications 14, 1217. 10.1038/s41467-023-36682-y.

9. Favre-Bulle, I. A., Vanwalleghem, G., Taylor, M. A., Rubinsztein-Dunlop, H., and Scott, E. K. (2018). Cellular-Resolution Imaging of Vestibular Processing across the Larval Zebrafish Brain. Current Biology 28, 3711–3722.e3. 10.1016/j.cub.2018.09.060.

10. Beiza-Canelo, N. et al. (2023). Magnetic actuation of otoliths allows behavioral and brain-wide neuronal exploration of vestibulo-motor processing in larval zebrafish. Current Biology 33, 2438–2448.e6. 10.1016/j.cub.2023.05.026.

11. Auer, F., Nardone, K., Matsuda, K., Hibi, M., and Schoppik, D. (2023). Purkinje Cells Control Posture in Larval Zebrafish (Danio rerio). bioRxiv. 10.1101/2023.09.12.557469.

12. Ehrlich, D. E. and Schoppik, D. (2017). Control of Movement Initiation Underlies the Development of Balance. Current biology : CB 27. du, 334–344. 10.1016/j.cub.2016.12.003.

13. Migault, G. and Bormuth, V. (2025). Light-sheet unit with two mirrors. Zenodo. 10.5281/zenodo.15498761.

14. 1. Randlett, O. et al. (2015a). Whole-brain activity mapping onto a zebrafish brain atlas. Nature methods 12, 1039–1046. 10.1038/nmeth.3581.

15. Greaney, M. R., Privorotskiy, A. E., D’Elia, K. P., and Schoppik, D. (2017). Extraocular motoneuron pools develop along a dorsoventral axis in zebrafish, Danio rerio. Journal of Comparative Neurology 525, 65–78. 10.1002/cne.24042.

16. Marquart, G. D., Tabor, K. M., Brown, M., Strykowski, J. L., Varshney, G. K., LaFave, M. C., Mueller, T., Burgess, S. M., Higashijima, S.-i., and Burgess, H. A. (2015). A 3D Searchable Database of Transgenic Zebrafish Gal4 and Cre Lines for Functional Neuroanatomy Studies. Frontiers in neural circuits 9, 1566. 10.1093/nar/gkr1288.

17. Muto, A., Ohkura, M., Abe, G., Nakai, J., and Kawakami, K. (2013). Real-Time Visualization of Neuronal Activity during Perception. Current Biology 23, 307–311. 10.1016/j.cub.2012.12.040.

18. Glasauer, S., Stephan, T., Kalla, R., Marti, S., and Straumann, D. (2009). Up–Down Asymmetry of Cerebellar Activation During Vertical Pursuit Eye Movements. The Cerebellum 8, 385–388. 10.1007/s12311-009-0109-5.

19. Stone, L. S. and Lisberger, S. G. (1990). Visual responses of Purkinje cells in the cerebellar flocculus during smooth-pursuit eye movements in monkeys. I. Simple spikes. Journal of Neurophysiology 63, 1241–1261. 10.1152/jn.1990.63.5.1241.

20. Krauzlis, R. J. and Lisberger, S. G. (1996). Directional organization of eye movement and visual signals in the floccular lobe of the monkey cerebellum. Experimental Brain Research 109, 289–302. 10.1007/bf00231788.

21. Kheradmand, A. and Zee, D. S. (2011). Cerebellum and Ocular Motor Control. Frontiers in Neurology 2, 53. 10.3389/fneur.2011.00053.

22. Ehrlich, D. E. and Schoppik, D. (2019). A primal role for the vestibular sense in the development of coordinated locomotion. eLife 8, e45839. 10.7554/elife.45839.

23. Iwamoto, Y., Kitama, T., and Yoshida, K. (1990). Vertical eye movement-related secondary vestibular neurons ascending in medial longitudinal fasciculus in cat I. Firing properties and projection pathways. Journal of Neurophysiology 63, 902–917. 10.1152/jn.1990.63.4.902.

24. Schoppik, D., Bianco, I. H., Prober, D. A., Douglass, A. D., Robson, D. N., Li, J. M. B., Greenwood, J. S. F., Soucy, E., Engert, F., and Schier, A. F. (2017). Gaze-Stabilizing Central Vestibular Neurons Project Asymmetrically to Extraocular Motoneuron Pools. The Journal of neuroscience : the official journal of the Society for Neuroscience 37, 11353–11365. 10.1523/jneurosci.1711-17.2017.

25. Goldblatt, D. et al. (2023). Neuronal birthdate reveals topography in a vestibular brainstem circuit for gaze stabilization. Current Biology 33, 1265–1281.e7. 10.1016/j.cub.2023.02.048.

26. Liu, Z., Hildebrand, D. G. C., Morgan, J. L., Jia, Y., Slimmon, N., and Bagnall, M. W. (2022). Organization of the gravity-sensing system in zebrafish. Nature Communications 13, 5060. 10.1038/s41467-022-32824-w.

27. Ahrens, M. B., Orger, M. B., Robson, D. N., Li, J. M., and Keller, P. J. (2013). Whole-brain functional imaging at cellular resolution using light-sheet microscopy. Nature methods 10, 413–420. 10.1038/nmeth.2434.

28. Wolf, S. et al. (2017). Sensorimotor computation underlying phototaxis in zebrafish. Nature Communications 8, 651. 10.1038/s41467-017-00310-3.

29. Dunn, T. W., Mu, Y., Narayan, S., Randlett, O., Naumann, E. A., Yang, C.-T., Schier, A. F., Freeman, J., Engert, F., and Ahrens, M. B. (2016). Brain-wide mapping of neural activity controlling zebrafish exploratory locomotion. eLife 5, e12741. 10.7554/elife.12741.

30. Bahl, A. and Engert, F. (2020). Neural circuits for evidence accumulation and decision making in larval zebrafish. Nature Neuroscience 23, 94–102. 10.1038/s41593-019-0534-9.

31. Dragomir, E. I., Štih, V., and Portugues, R. (2019). Evidence accumulation during a sensorimotor decision task revealed by whole-brain imaging. Nature neuroscience 23, 85–93. 10.1038/s41593-019-0535-8.

32. Poulsen, R. E., Scholz, L. A., Constantin, L., Favre-Bulle, I., Vanwalleghem, G. C., and Scott, E. K. (2021). Broad frequency sensitivity and complex neural coding in the larval zebrafish auditory system. Current Biology 31, 1977–1987.e4. 10.1016/j.cub.2021.01.103.

33. Favre-Bulle, I. A., Taylor, M. A., Marquez-Legorreta, E., Vanwalleghem, G., Poulsen, R. E., Rubinsztein-Dunlop, H., and Scott, E. K. (2020). Sound generation in zebrafish with Bio-Opto-Acoustics. Nature Communications 11, 6120. 10.1038/s41467-020-19982-5.

34. Balaban, C. (2016). Chapter 3 Neurotransmitters in the vestibular system. Handbook of Clinical Neurology 137, 41–55. 10.1016/b978-0-444-63437-5.00003-0.

35. Vladimirov, N., Mu, Y., Kawashima, T., Bennett, D. V., Yang, C.-T., Looger, L. L., Keller, P. J., Freeman, J., and Ahrens, M. B. (2014). Light-sheet functional imaging in fictively behaving zebrafish. Nature methods 11, 883–884. 10.1038/nmeth.3040.

36. Montgomery, D. C. and Runger, G. C. (2011). Applied Statistics and Probability for Engineers, 2th Edition. Wiley Global Education. Wiley Global Education.

37. Rohlfing, T. and Maurer, C. R. (2003). Nonrigid image registration in shared-memory multiprocessor environments with application to brains, breasts, and bees. IEEE trans. inf. technol. biomed. 7, 16–25. 10.1109/TITB.2003.808506.

38. Randlett, O. et al. (2015b). Whole-brain activity mapping onto a zebrafish brain atlas. Nat. Methods 12, 1039–1046. 10.1038/nmeth.3581.

39. Migault, G., Beiza-Canelo, N., Chatterjee, S., Debrégeas, G., and Bormuth, V. (2026). Numerical source data for graphs and charts for : Directionally Biased Neuronal Responses to Pitch-Axis Vestibular Stimulation in Larval Zebrafish Compared to Roll-Axis Responses. Zenodo. 10.5281/zenodo.19253364.

